# Age-related Decline of a Chaperone Complex in the Mouse Heart

**DOI:** 10.1101/2024.09.12.612759

**Authors:** Purnima Devaki Bhat, Aswan Nalli, Luke Shoemaker, Jennifer Q. Kwong, Hiroshi Qadota, Prasanna Krishnamurthy, Guy M. Benian

## Abstract

The exquisitely organized sarcomere, the unit of contraction of striated muscle, is a stable structure with slow turnover of its components. The myosin chaperone UNC-45 and its binding partners, Hsp90 and Hsp70, are required for the initial folding of the myosin head domain and the assembly of myosin into thick filaments. There is increasing evidence that the UNC-45 system has an important role during aging to preserve sarcomere organization. Its decline may be a key factor in sarcopenia. Unlike skeletal muscle, the UNC-45 system in cardiac muscle in aging heart has not been examined extensively. Here we show that Unc45b and Hsp70 are localized to sarcomeric Z-discs in the mouse heart. We further show that during aging, there is a decline in the levels of myosin heavy chain, Unc-45b and Hsp70, but not Hsp90. While the decrease in Unc45b appears to be at the mRNA level, the decrease in the levels of myosin and Hsp70 were not at the mRNA but at the protein level. We have reported that in skeletal muscle, there is a decline in both Unc45b and Hsp90, and here we show that there is no such decline of Hsp70 in skeletal muscle. Hsp70 levels also did not decline with age in the brain or the liver. This heart-specific decrease of Hsp70 through its function as an Unc45b/Hsp70 complex might account for the age-dependent worsening of cardiomyopathies, and through Hsp70’s multiple Unc-45b-independent functions, affect the folding and assembly of many other proteins in the aging heart.

## 1. INTRODUCTION

Muscle contraction results from the pulling inwards of thin filaments anchored to Z-lines by myosin head domains located on the surface of thick filaments. Myosin is a hexamer consisting of two myosin heavy chains, and four myosin light chains, and has two head domains, two necks, and a rod that is comprised of the two myosin heavy chains in an α-helical coiled coil structure. The myosin rods assemble into the thick filament shaft with the myosin heads displayed on their surface in a specific pattern. The myosin heads are nanometer scale motors that efficiently convert the energy of ATP hydrolysis, through a series of conformational changes, to the swinging of the neck of myosin while a single head is attached to a thin filament.

UNC-45 is a chaperone required to fold the myosin head after translation and for assembly of myosin into thick filaments during development (Barral et al., 1998, 2002). UNC-45 may also be involved in refolding the myosin head after stress in adulthood (Etard et al., 2008; Kachur and Pilgrim, 2008). A role for UNC-45 in mature muscle is plausible given that there is a relatively slow turnover of myosin in assembled thick filaments (Solomon and Goldberg, 1996). Recently, we have reported that UNC-45 has an important role during adult aging to preserve myosin levels and myofibril organization in body wall muscle, the major striated muscle in *C. elegans* (Matheny et al., 2024). Moreover, we found that additional expression of UNC-45 in old adults leads to maintenance of myosin levels and prevents the disorganization of myofibrils, suggesting that increasing UNC-45 might be a way to prevent or treat sarcopenia, the age dependent decline in sarcomere organization and function in skeletal muscle that occurs without an underlying condition.

UNC-45 was first discovered by genetic analysis in *C. elegans*, by finding a temperature sensitive mutant that when grown at the restrictive temperature results in an adult worm with highly disorganized myofibrils and reduced muscle function (Epstein and Thomson, 1974). It was later found that *unc-45* null alleles are embryonic lethal (Venolia and Waterston, 1990; Barral et al., 1998). Whereas *C. elegans* and Drosophila have single Unc-45 genes, humans and mice have two genes for UNC-45, one encoding Unc-45b, expressed in striated (skeletal and cardiac) muscle cells, and the other, Unc-45a, expressed in all cell types (Price et al., 2002).

All UNC-45 homologs consist of a C-terminal UCS domain which binds to the myosin head, an N-terminal TPR domain that binds to HSP-90 (Barral et al, 2002) and HSP-70 (Gazda et al., 2013), and a central region that acts as an inhibitor of the myosin power stroke (Bujalowski et al., 2018). Evidence that UNC-45 acts as a chaperone for the myosin head includes showing that: (1) UNC-45 prevents the thermal aggregation of myosin heads (Barral et al., 2002); (2) through a clever single molecule experiment that UNC-45 can refold an unfolded myosin head (Kaiser et al., 2012); and (3) UNC-45 can protect the myosin head’s Mg-ATPase activity from being reduced by heat denaturation (Gaziova et al., 2020). The crystal structure of nematode UNC-45 shows that it forms linear multimers, two to five subunits each, in which the repeating unit spans 17 nm, which is similar to the absolutely conserved repeating unit of 14.3 nm of the staggered display of pairs of myosin heads along the surface of thick filaments (Gazda et al., 2013). Therefore, such UNC-45 multimers could help direct or stabilize this arrangement of myosin heads during thick filament assembly. In addition, this architecture suggests the possibility that in the assembled thick filament, refolding of myosin heads that were damaged by oxidative or thermal stresses as muscle is used and ages, might be facilitated. Support for such a role in mature muscle is that in at least *C. elegans*, antibodies to UNC-45 co-localize with myosin on the major portions of the thick filaments in already assembled sarcomeres of adult muscle (Ao and Pilgrim, 2000; Gazda et al., 2013). In zebrafish skeletal muscle, Unc-45b normally localizes to Z-disks, but moves to thick filaments during stress (cold and heat shock, chemical and physical stress)(Etard et al., 2008).

UNC-45 has also been reported to have crucial functions in heart muscle. RNAi-mediated knockdown of *unc-45* in the single-tubed heart of Drosophila results in reduced myofibrils, myofibril disorganization, decreased myosin accumulation, reduced cardiac contractility, dilation and arrhythimias, especially if the RNAi is conducted during morphogenesis (Melkani et al. 2011). Surprisingly, Unc-45b may also function as a chaperone for the cardiogenic transcription factor GATA4 (Chen et al., 2012). Unc-45b forms a physical complex with GATA4, although chaperone activity has not been demonstrated. Interestingly, recessive loss of function mutations in the Unc-45b gene in mice result in reduced levels of GATA4 (in addition to reduced levels of myosin) and arrest of cardiac morphogenesis and embryonic lethality.

Like humans, *C. elegans* develops sarcopenia, a reduction in the number and organization of sarcomeres and skeletal muscle function with age (Herndon et al., 2002; Matheny et al., 2024). We have reported that using a temperature sensitive *unc-45* mutant to reduce the level and function of UNC-45 at the beginning of adulthood, sarcopenia develops earlier. We also observed a sequential decline of HSP-90 (adult day 3), UNC-45 (day 4) and then myosin (day 8), perhaps reflecting a cause and effect relationship (Matheny et al., 2024). The decline of these proteins is likely to be the result of protein degradation as the decline in their mRNAs occurs much earlier, especially for myosin (mRNA at day 1, protein at day 8). We have also shown that in mouse skeletal muscle Hsp90 and Unc-45b are reduced when comparing 3 month old to 24 month old mice (Matheny et al., 2024). No such studies have been undertaken to examine the Unc45 chaperone system in the aging heart, and a process similar to sarcopenia could play an important role in cardiac dysfunction with age.

Here we report our examination of the UNC-45 chaperone complex in the aging mouse heart. We find that when comparing 3 month old to 22 month old hearts, there is a decline in myosin (myosin heavy chains), Unc-45b and Hsp70, but not Hsp90, or GATA4 or OLA1. The decline in Hsp70 in the heart was particularly dramatic, and seems to be heart-specific since we did not observe an age-dependent decline in Hsp70 in skeletal muscle, liver and brain. Interestingly, while the decrease in Unc45b could be at the mRNA level, the decrease in Hsp70 appears to be at the protein and not at the mRNA level. Although Hsp70 forms a complex with Unc45b, the role of Hsp70 in the heart is likely to extend beyond its cooperation with Unc45b to initially fold myosin and refold damaged myosin.

Cardiac aging at the cellular level consists of multiple changes including cardiomyocyte hypertrophy, increased cardiomyocyte apoptosis, increased numbers of fibroblasts, and increased collagen deposition (Vijayakumar et al. 2024). However, the molecular mechanisms for these alterations is poorly understood. Cardiomyocyte hypertrophy is thought to be a compensatory mechanism for the loss of cardiomyocyte numbers and consists of an increase in cell size, increase in protein synthesis, changes in sarcomere organization (Frey et al. 2004), and increased numbers of myofibrils in surviving cardiomyocytes. The results described here provide further molecular information about what happens during aging in the heart.

## 2. RESULTS

### 2.1. Unc45b and Hsp70 are localized to the Z-discs of cardiomyocytes

Unc45 forms a multimeric chain (Figure 1A), which then complexes with Hsp90 and Hsp70 (see ref. Gazda et al., 2013). This complex interacts with the motor domains of newly translated myosin heavy chains to assure their proper folding, and is required for myosin to be assembled into thick filaments, and also probably refolds damaged mature myosin (Lee et al., 2014). Since there is no information where Unc45b and Hsp70 are actually localized within cardiomyocytes, we immunostained heart sections from 3-month and 24-month old C57 Black mice. As shown in Figure 1B, C, both Unc45b and Hsp70 are co-localized with α-actinin at Z-discs of cardiomyocytes. These results are consistent with the fact that Unc45b and Hsp70 form a complex. The localization of Unc45b to Z-discs in mouse cardiomyocytes is similar to the localization of Unc45 to Z-discs of zebrafish skeletal muscle (Etard et al., 2008). Our immunostaining did not reveal any obvious changes in the localization or abundance of Unc45b or Hsp70 in young vs. old mouse hearts.

**Figure 1:**
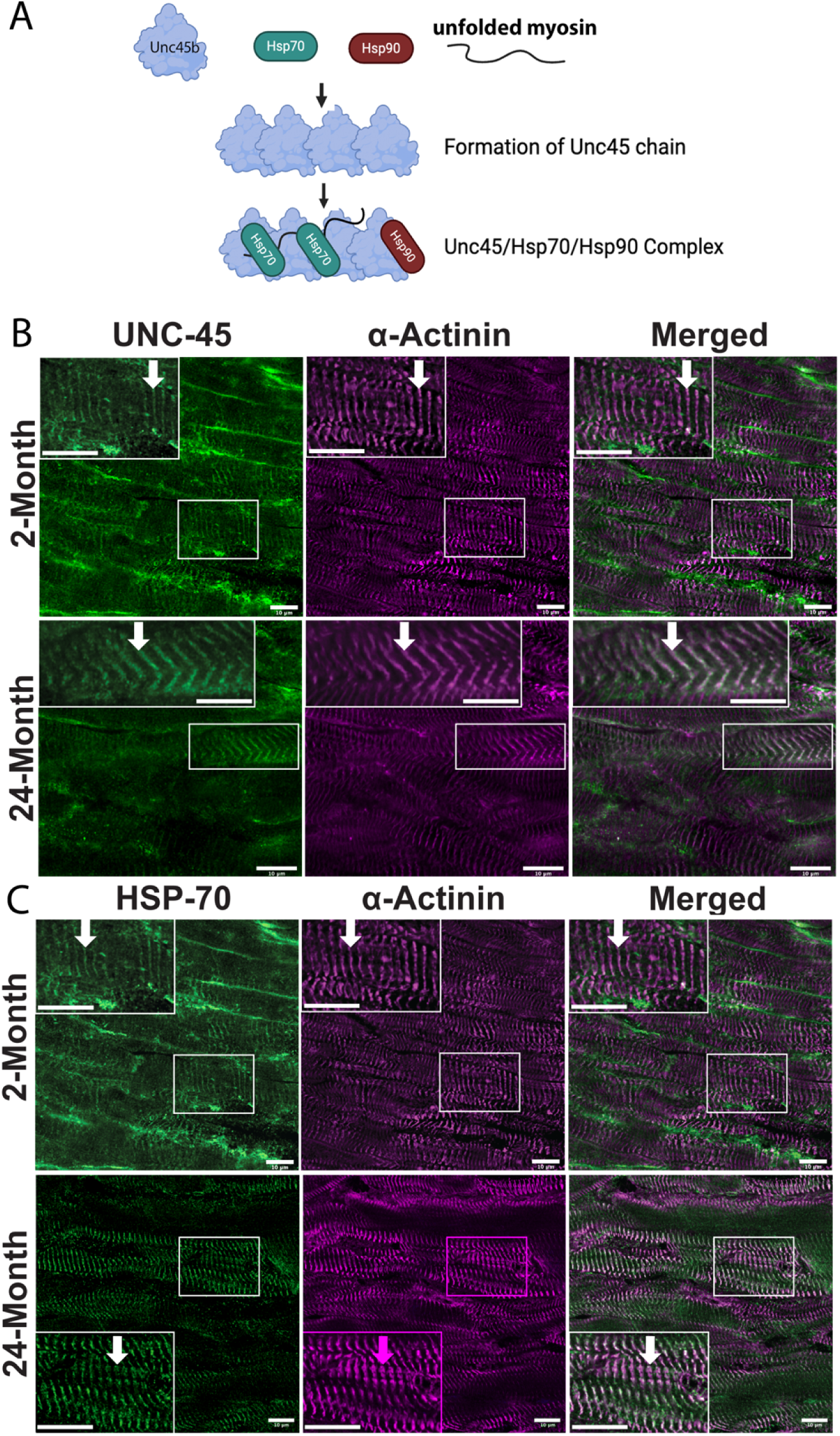
Unc45b, Hsp70 and Hsp90 work together to fold the motor domain of myosin. **(A):** The crystal structure of the *C. elegans* UNC-45/HSP-70/HSP-90 complex (Gazda et al. 2013) shows that UNC-45 (or by extension, mouse Unc45b) forms linear multimers and recruits Hsp70 and Hsp90, and each of the Unc45b/Hsp70 or Unc45b/Hsp90 complexes folds one myosin head. The periodicity of these complexes matches the periodicity of the spacing of myosin heads on the surface of the assembling thick filament. **(B and C): Unc45b and Hsp70 are localized to sarcomeric Z-discs in cardiomyocytes.** Heart sections from 3 and 24-month old C57BL/6J mice were co-immunostained with anti-Unc45b and anti-α-actinin (panel B), or anti-Hsp70 and anti-α-actinin (panel C). α-actinin marks the Z-discs. Arrows on the insets point to single Z-discs. The scale bar, 20 μM.

### 2.2. Myosin, Unc-45b, and Hsp70, but not GATA4, Hsp90, and OLA1 are reduced in aged mouse hearts

We harvested hearts from C57 Black mice at 3 months and 22 months of age, which translates roughly to 20 years and 65 years in humans (Dutta and Sengupta, 2016), and prepared total protein lysates. These lysates were used to perform quantitative western blots to compare the levels of several proteins known to be part of an Unc-45b chaperone complex (Figure 1A). As shown in Figure 2A, we observed an approximately 30% decrease in the level of the UNC-45b client, myosin heavy chains, in 22 month old vs. 3 month old hearts. In contrast, we observed no change in the level of the possible Unc-45b client protein GATA4 (Figure 2B). Next, we examined the levels of Unc-45b, and found approximately a 25% decline in the level of Unc-45b in both male and female mice (Figure 3A). Interestingly, a dramatic (∼60%) decline in the level of Hsp70 was observed between 3 month and 22 month old mouse hearts (Figure 3B), while no significant decline was seen with Hsp90 levels between the two ages (Figure 3C). Similarly, there was no downregulation of another protein called OLA1 or Obg-like ATPase 1 (Figure 3D), an ATPase protein involved in cardiomyopathies (Dubey et al., 2024).

**Figure 2:**
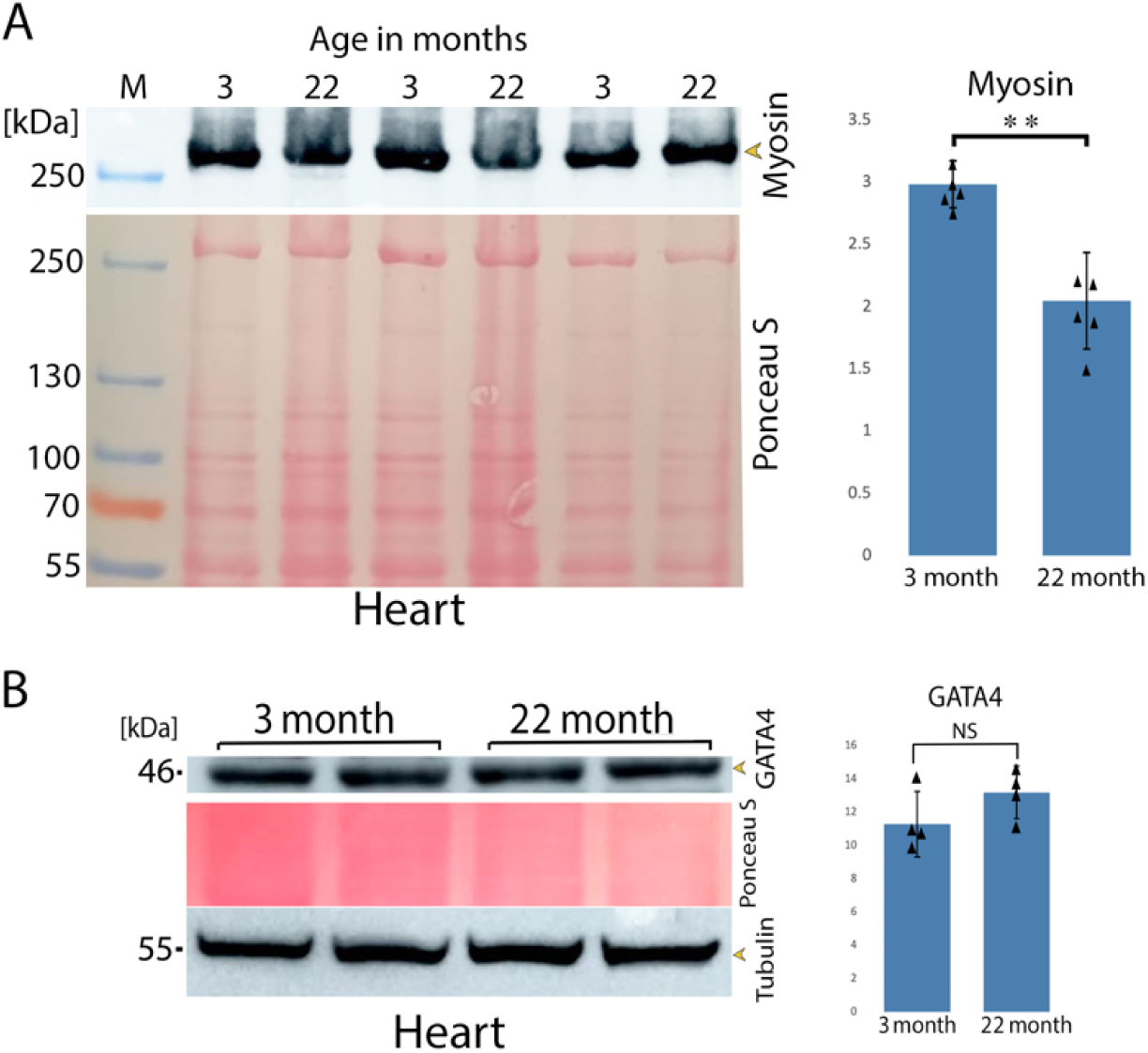
Myosin but not GATA4 levels are reduced in aged mice. **Panels A and B:** Western analysis of hearts from 3 months and 22 months old mice using an antibodies against myosin heavy chain (A) or GATA4 (B). Ponceau S was used for loading control for myosin and both Ponceau S and Tubulin were used as control for GATA4. The results were subjected to statistical analysis using a student’s t test.**: P<0.01; NS: not significant.

**Figure 3:**
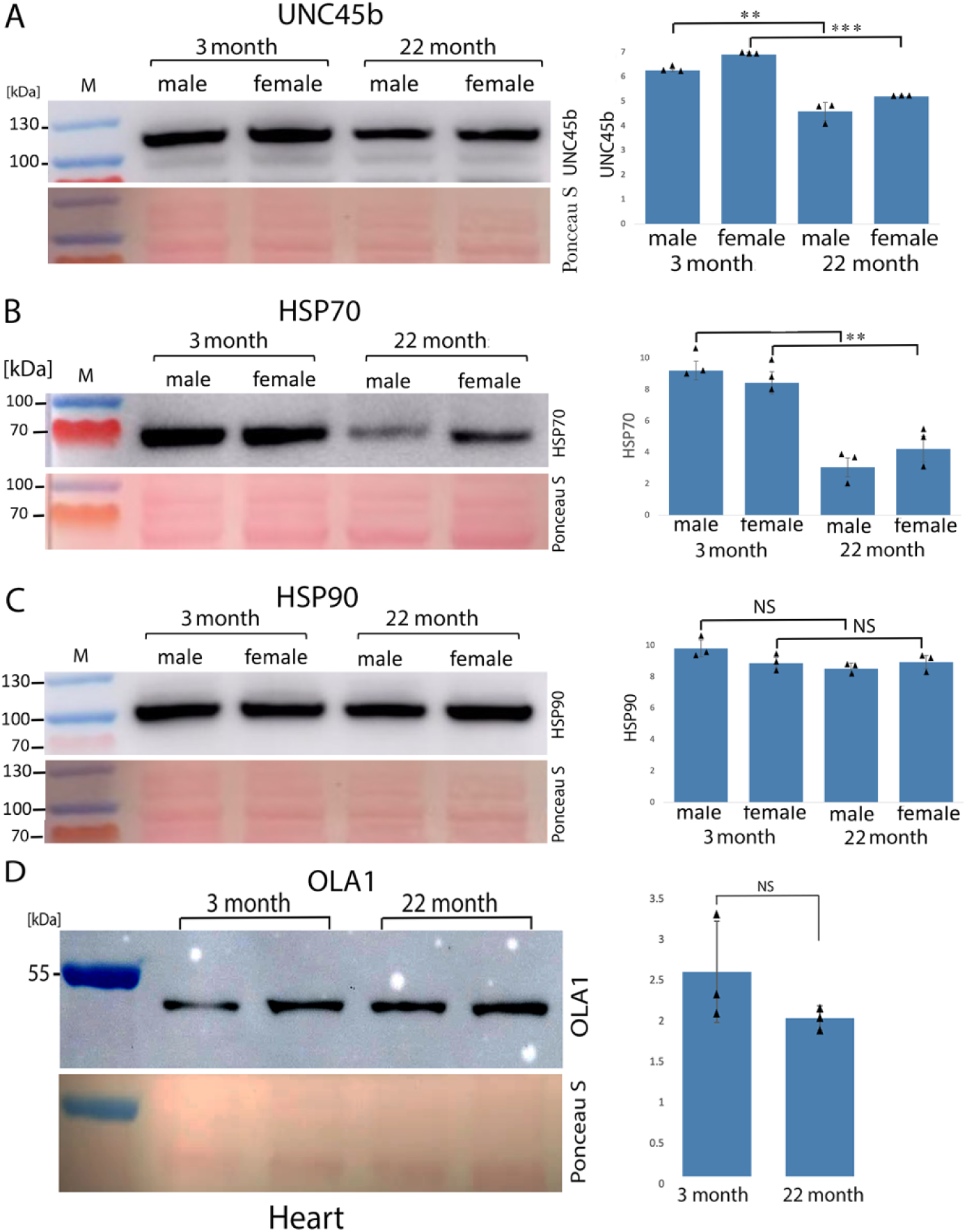
UNC45b and HSP70 but not HSP90 or OLA1 are reduced in aged mouse hearts. Western analysis of hearts from 3 months and 22 month old male and female mice using antibodies against Unc45b (A), Hsp70 (B), and Hsp90 (C), or OLA1 (D). Ponceau S was used for loading control. The results were subjected to statistical analysis using a student’s t test. **: P<0.01; ***: P<0.001; NS: not significant.

### 2.3. Downregulation of Hsp70 with age is specific to the heart

Previous results have shown that both Unc-45b and Hsp90 are downregulated in quadriceps skeletal muscle with age (Matheny et al. 2024). To determine if Hsp70 is also downregulated in skeletal muscle, we made samples from 3 month and 22 month old quadriceps skeletal muscle and performed quantitative Western analysis. As shown in Figure 4A, no such decline in Hsp70 was observed with age in skeletal muscle. Additionally, we also examined Hsp70 levels in aged mice liver (Figure 4B) and brain (Figure 4C), and found no significant decline in the levels with age. These results indicate that the decline of Hsp70 levels with age is specific to the heart.

**Figure 4:**
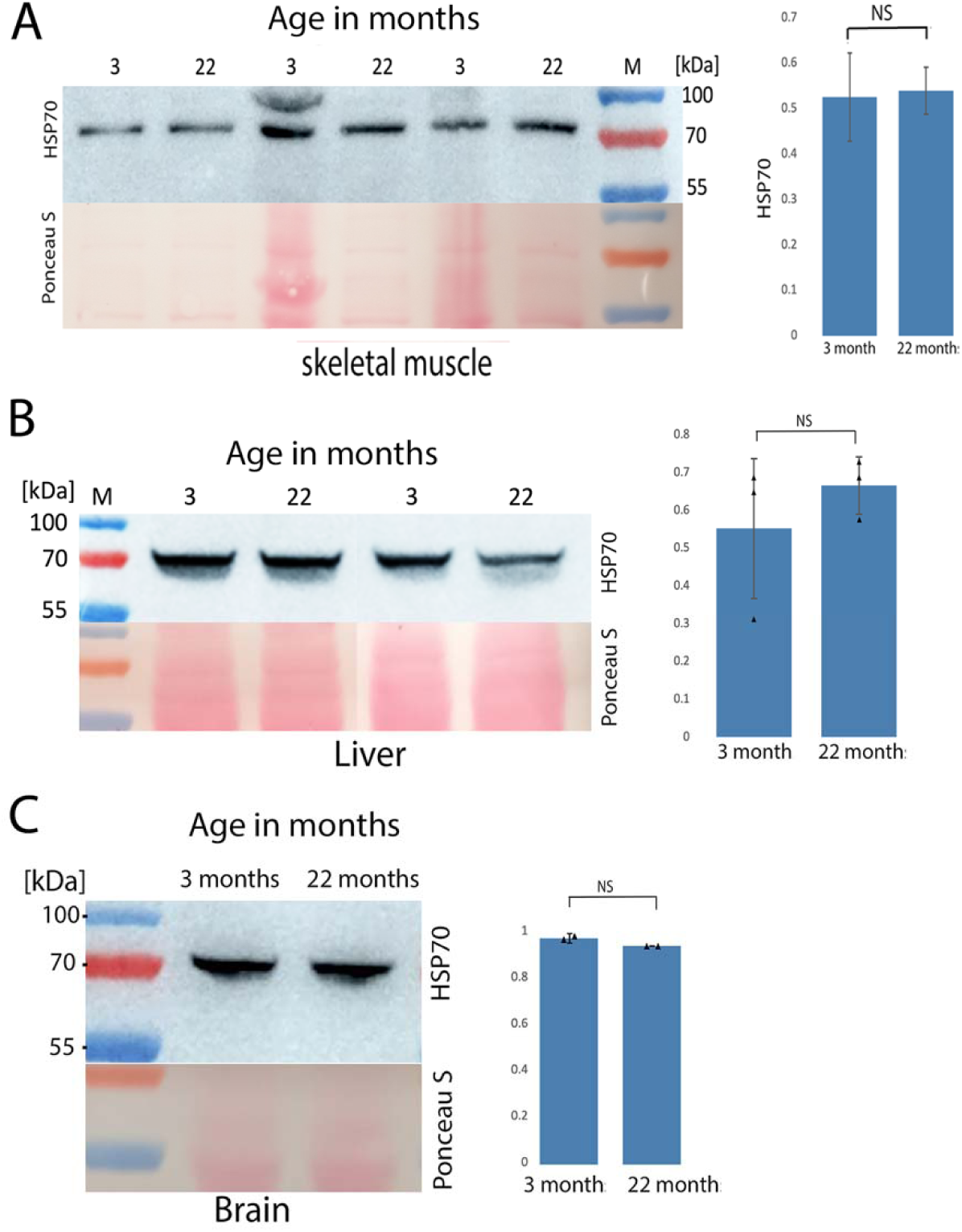
Hsp70 was not downregulated with age in skeletal muscle, liver or brain. Western analysis of skeletal muscle (A), liver (B) and brain (C) from 3 month- and 22 month-old mice using an antibody Hsp70. Ponceau S staining was used for loading control. The results were subjected to statistical analysis using a student’s t test. NS: not significant.

### 2.4. The downregulation of Unc-45b but not Hsp70 occurs at the mRNA level

We next examined if the downregulation of myosin, Unc-45b and Hsp70 occurs at the transcriptional level using RT-qPCR. RNA samples were prepared from the 3 month and 22 month old mouse hearts, reverse-transcribed and subjected to qPCR. As shown in Figure 5, we found that Unc-45b downregulation with age appears to occur at the mRNA level since the difference between the two ages was statistically significant (Figure 5B). Myosin heavy chain mRNA also appears to decline with age, although not reaching statistical significance (Figure 5A). However, there was no difference in the levels of mRNA for Hsp70 between 3 and 22 months of age (Figure 5C), indicating that the Hsp70 downregulation occurs at the protein level. There was also no difference in the mRNA levels for Hsp90 between the two ages (Figure 5D).

**Figure 5:**
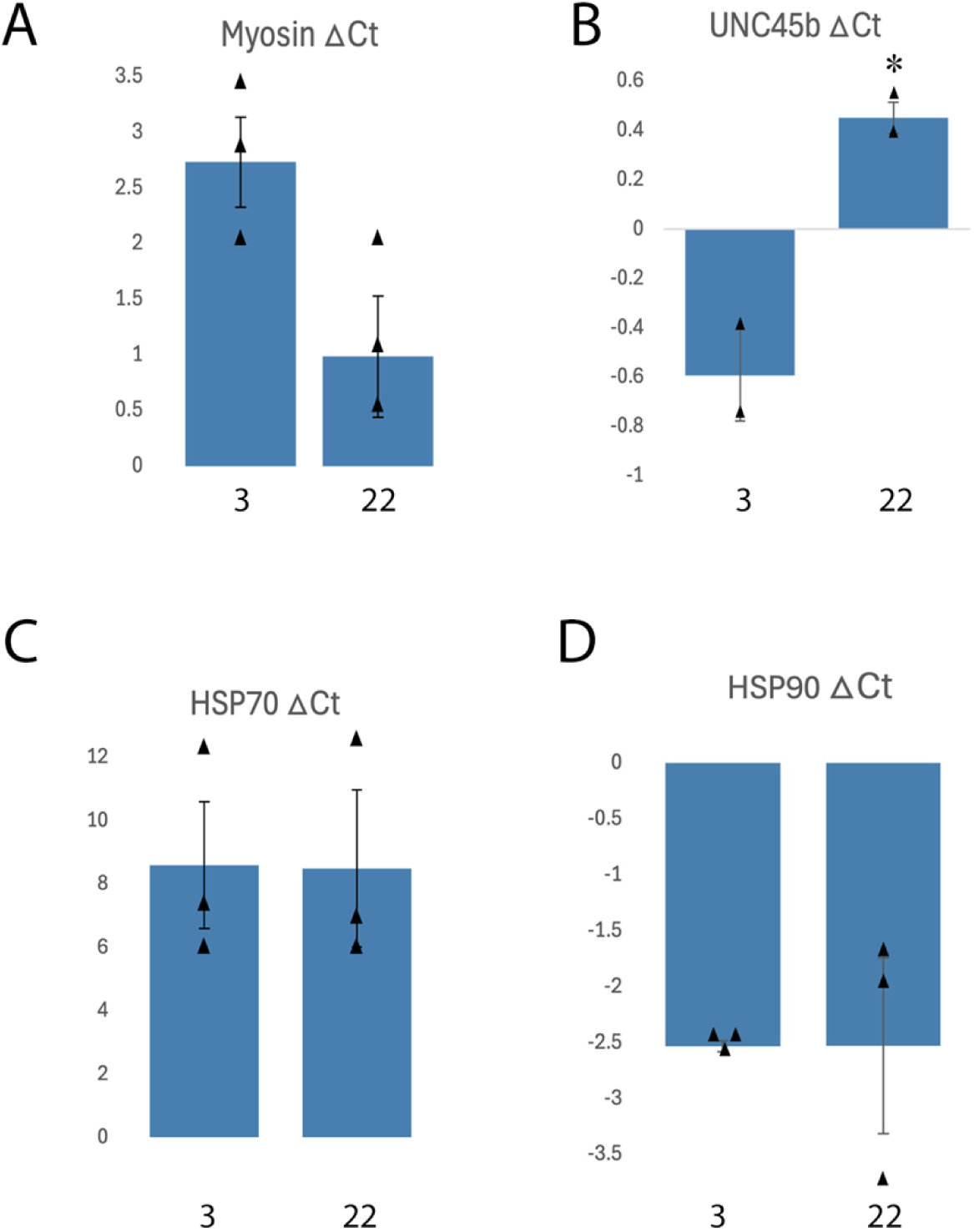
The downregulation of UNC-45b and myosin but not Hsp70 or Hsp90 occurs at the mRNA level. Analysis of mRNAs for myosin heavy chain, UNC45b, Hsp70 and Hsp90 by RT-qPCR. The mRNA were isolated from 3 different hearts, aged 3 and 22-months, reverse transcribed and subjected to qPCR. The ΔCt values were calculated and plotted as histograms. ΔCt value is negative whenever the value for the gene of interest is higher than the housekeeping gene (GAPDH). Thus, a lower-expressed gene will be positive. qPCR detects higher expression at a lower Ct and low expression at higher Ct.

### 2.5. Cardiomyocyte organization is disrupted with age in mice hearts

Because skeletal muscle sarcopenia includes a reduction in myofibrils and sarcomeric disorganization, we made a preliminary analysis of muscle cardiomyocyte organization in young vs. old hearts. We used antibodies to to α-actinin to visualize Z-discs, and antibodies to N-cadherin to visualize intercalated discs. Our α-actinin staining reveals that the cardiomyocytes in the 22 month old hearts are less regularly organized—instead of tight parallel packing there are loose angled arrays (Figure 6A). N-cadherin staining reveals a greater variation in intercalated disk shapes and sizes with overall lower intensity staining (Figure 6B).

**Figure 6:**
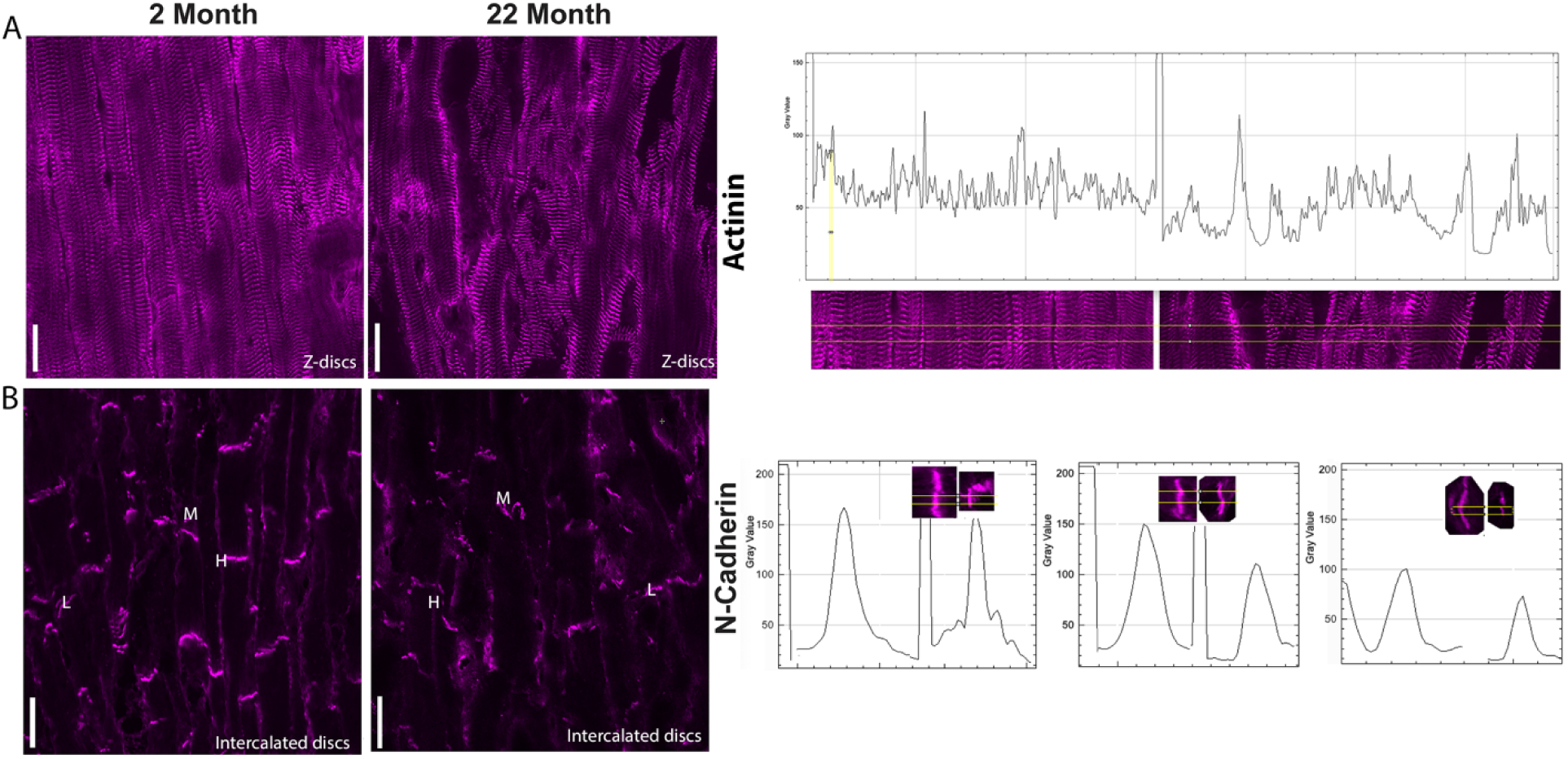
Disorganization of cardiomyocytes within the myocardium and greater variation in intercalated disc shapes and sizes with age. The organization of cardiomyocytes within the myocardium was examined in young vs. old hearts by immunohistochemistry. Immunostaining was done with antibodies to α-actinin for Z-disks (A) and against N-Cadherin for intercalated discs (B). Longitudinal views are shown. The images were also analyzed with ImageJ software using the Plot profile. Scale bar, 20 μm.

## 3. DISCUSSION

Here, we show that the Unc45b chaperone system is significantly downregulated with age in mouse hearts. The levels of myosin, Unc45b, and Hsp70 declined between 3 months and 22 months in mouse hearts. In contrast, we found no decreases in the levels of Hsp90, GATA4 and the unrelated protein, OLA1. The reduction does not appear to be at the transcriptional level. Moreover, we found that Hsp70 downregulation was specific to the heart, as we found no differences in the levels of Hsp70 between 3 and 22 month old mouse liver, brain, or skeletal muscle. In Matheny et al. (2024) we primarily reported on sarcopenia in *C. elegans*, but also showed that there is approximately a 60% decline in both Unc45b and Hsp-90 levels between 3 month and 24 month old mouse quadriceps skeletal muscle. Taken together with our present study, the following picture emerges. In mice, for both skeletal and heart muscle, there is an age dependent decrease in Unc45b and myosin, but in skeletal muscle there is also a decline in Hsp90, and in cardiac muscle there is also a decline in Hsp70. Thus, depending on the type of striated muscle, a different heat shock protein that binds to Unc45b undergoes an age-dependent decline.

It is not clear what the functions of Hsp90 and Hsp70 are in the Unc45 complex. From biochemical and structural studies of the UNC-45/HSP-90/HSP-70 complex in *C. elegans*, the same binding site of the N-terminal TPR domain of UNC-45 binds to either HSP-90 or HSP-70, with the binding to HSP-70 having less affinity (KD of 120 μM for HSP-70 vs. KD of 14 μM for HSP-90)(Gazda et al. 2013). These authors also suggest that HSP-70 and HSP-90 may either occupy alternating positions along the UNC-45 oligomer or bind sequentially to carry out early vs. late folding steps. However, it is not clear what the function of HSP-90 and HSP-70 are in the UNC-45 complex. Given that UNC-45 by itself has been shown to have chaperone activity on myosin (Barral et al., 2002; Kaiser et al., 2012; Gaziova et al., 2020), one possibility is that HSP-90 and HSP-70 provide additional chaperoning activities for myosin, and the binding of UNC-45 to myosin positions the complex so that HSP-90 and HSP-70 can provide this activity. An alternative possibility is that the binding of HSP-90 and HSP-70 to UNC-45 simply stabilizes UNC-45.

We showed that both Unc45b and Hsp70 are located at Z-discs, in both young and old hearts (Figure 1B, C). This result seems surprising given that the function of Unc45/Hsp70 and Unc45/Hsp90 complexes are to fold myosin upon translation, or in the case of already mature muscle, to refold myosin heads which are located on the surface of thick filaments which are located at A-bands. However, there is precedence for UNC-45 normally being located at Z-disks in the skeletal muscle of zebrafish (Etard et al. 2008). In this muscle type, UNC-45 re-localizes to A-bands upon thermal or ROS stress (Etard et al. 2008), presumably to either refold damaged myosin heads, or ensure proper folding of newly synthesized myosin that replaces damaged myosin. Thus, in this and our context, the Z-disc acts as a “reservoir” for Unc45b and Hsp70, and these proteins can be mobilized, by some unknown mechanism, to where they are needed at the A-band.

Certainly, even in a muscle cell, Hsp90 and Hsp70 each have many functions independent of UNC-45. Much of Hsp70 may be in the cytosol and not detectable by our immunostaining methods. Hsp70 is likely to fold many proteins in cardiomyocytes, and a decline in Hsp70 could affect multiple processes in cardiomyocytes, not just the level and or organization of myofibrils. Moreover, our results need to be interpreted with some caution, as the extracts we have made are from whole hearts and the heart is cellularly heterogeneous.

Normally, Hsp70 binds to polypeptides as they are being synthesized to prevent them from aggregating. Once the client protein is fully synthesized, Hsp70 can transfer the client protein to other chaperones. Proteins can also be transferred from Hsp70 to Hsp90 by the Hsp70/Hsp90 Organizing Protein (Hop/Stip1/Sti1)(Bhattacharya et al. 2020) and is indeed expressed in cardiomyocytes according to The Human Protein Atlas.

In addition, in the heart, although cardiomyocytes are the prominent cell type, there are also fibroblasts, endothelial cells, smooth muscle cells and even inflammatory cells. In fact, in the young heart 65-80% of the total heart volume is from cardiomyocytes, but during normal aging up to 30% of these are lost from apoptosis (Sheydina et al. 2011), and there is no regeneration. We think the decline of both Unc45b and Hsp70 levels with age is unlikely due to this loss of cardiomyocytes since the levels of GATA4, and Hsp90 were not downregulated (see Figure 2B and Figure 3C). Similarly, there was no downregulation of another protein called Ola1 or Obg-like ATPase 1 (Figure 3D), an ATPase protein involved in cardiomyopathies (Dubey et al., 2024). It is possible that no significant apoptosis occurs at 22 months of age, but later on.

While there may be a loss of cardiomyocytes with aging at some point, perhaps beyond 22 months of age, as a seemingly adaptive mechanism to maintain heart function, the remaining cardiomyocytes undergo hypertrophy (Vijayakumar et al., 2024) due to an increase in myofibrils per cell. The myofibrillogenesis which includes new myosin translation and new thick filaments certainly require Unc45 and Hsp70, thus our observation of significant declines in these proteins suggests that the newly synthesized myosin may not be optimally folded or optimally functional and thus contribute to a decline in contractile activity and heart pumping. Nonetheless, the finding that both Unc45b and Hsp70 are downregulated with age in the heart offers certain insight into cardiac dysfunction. Their reduced levels may play a role in the worsening of cardiomyopathy phenotype during aging, regardless of the causative mutations. Given that Hsp70 has a role in protein stabilization and preventing protein aggregation, its age-dependent reduction could potentially affect diseases such as cardiac amyloidosis. In cardiac amyloidosis, amyloid protein, which is misfolded and insoluble, aggregates and gets deposited in the cardiac muscle and other heart cells (Fikrle et al., 2012; Banypersad et al., 2012). These aggregate deposits cause irreversible heart thickening and heart disease. Thus, a reduction in the level of Hsp70 could contribute to the misfolding and aggregation of amyloid protein in cardiac amyloidosis.

Finally, the reduction in the levels of Hsp70 in 22 month old mice heart does not appear to be at the transcriptional level (Figure 5C). The reduction is unlikely due to a loss of ribosomal function with age since the levels of several other proteins examined such as GATA4 and OLA1 did not decline with age. We suspect two possibilities, one is that there is an age-dependent increase in microRNAs inhibiting translation of the Hsp70 mRNA, and or there is an age-dependent increase in ubiquitin-proteasomal degradation of Hsp70.

## 4. MATERIALS AND METHODS

### 4.1. Mice age and genotype

Young and old mice were of the genotype C57BL/6J were used. Young age is 2 or 3 months old and old is 22 or 24 months old. Mice were all healthy and hearts and skeletal muscle were harvested and samples were prepared as described below.

### 4.2. Western blot analysis

Heart ventricular tissues (1 cm³) from C57 black mice aged three months and 22 months were collected. The tissues were homogenized in chilled 650 µL RIPA buffer (50 mM Tris pH 7.5, 150 mM NaCl, 1 mM EDTA, 0.5% NP40, 0.1% SDS, and PIC [final concentration: 1x, Sigma Aldrich P2714]) using a Bead Ruptor Elite (Speed: 5 m/s, Time: 30 s, Cycles: 15), with the tissue kept on ice between cycles. The lysates were centrifuged at 13,000 rpm for 5 minutes in an Eppendorf microfuge. The supernatant (∼520 µL) was transferred to a clean 1.5 mL tube on ice and mixed with 200 µL of 4X Loading Dye and 80 µL of 10X Reducing Agent (Invitrogen NuPAGE NP007 and NP009, respectively). The samples were heated at 95°C for 5 minutes and stored at −20°C until use. Five µL of each sample were loaded per lane on a 4-12% Tris-acetate gel (Invitrogen XP04125BOX), and proteins were transferred to a 0.2 µm nitrocellulose membrane (Bio-Rad) using the Trans-Blot Turbo Transfer System (Bio-Rad). After transfer, membranes were stained with 25 mL Ponceau S Stain (Sigma Aldrich P7170) for 5 minutes, washed extensively with water, and imaged for loading control. Membranes were blocked with 5 mL of 1x TBST (Tris Buffered Saline with 0.1% Tween-20) containing 5% powdered milk for one hour at room temperature. After blocking, membranes were washed and incubated in 5 mL of primary antibody solution (primary antibody diluted in 1x TBST + 1% powdered milk) overnight at 4°C on a BenchRocker (Benchmark Scientific). The primary antibodies used were against UNC45B (rabbit, 1:2,500, Proteintech 21640-1-AP), HSP70 (rabbit, 1:7,000, Proteintech 10995-1-AP), HSP90 (rabbit, 1:5,000, Proteintech 13171-1-AP), and Myosin (mouse,1:1,000, Abcam ab50967).The primary antibody solutions were removed, and membranes were washed three times for 15 minutes each in 1x TBST. Following the washes, membranes were incubated with secondary antibody solutions diluted in 1x TBST + 1% powdered milk to a final volume of 5 mL. Secondary antibodies included HRP-conjugated Goat anti-Rabbit (1:15,000) and Goat anti-Mouse (1:100,000) (Jackson Laboratories), and were incubated for 2 hours at room temperature on a BenchBlotter 2D Rocker (Benchmark Scientific). Membranes were washed three times for 15 minutes each in 1x TBST to remove excess antibodies. Finally, membranes were treated with Immobilon Western Chemiluminescent HRP Substrate (MilliporeSigma) according to the manufacturer’s instructions and imaged using an Amersham Imager 600 (GE Healthcare Life Sciences).

### 4.3. RNA Extraction and Real-Time Quantitative PCR

Mouse heart tissue (∼1 cm³) was homogenized in RNA lysis buffer (RLT) using a Bead Ruptor Elite (6 m/s, 30 s, 2 cycles; 2.8 mm ceramic beads), following the protocol provided in the Qiagen RNeasy Mini Kit (Cat. No. 74104). Total RNA was extracted per the manufacturer’s instructions. For qRT-PCR, 150 ng of RNA was used as a template with the Luna Universal One-Step RT-qPCR Kit (Cat. No. E3005S), according to the manufacturer’s protocol, on the Thermo Fisher QuantStudio 5 system. The nucleotide sequences for the oligos used in the study are provided in the table below.

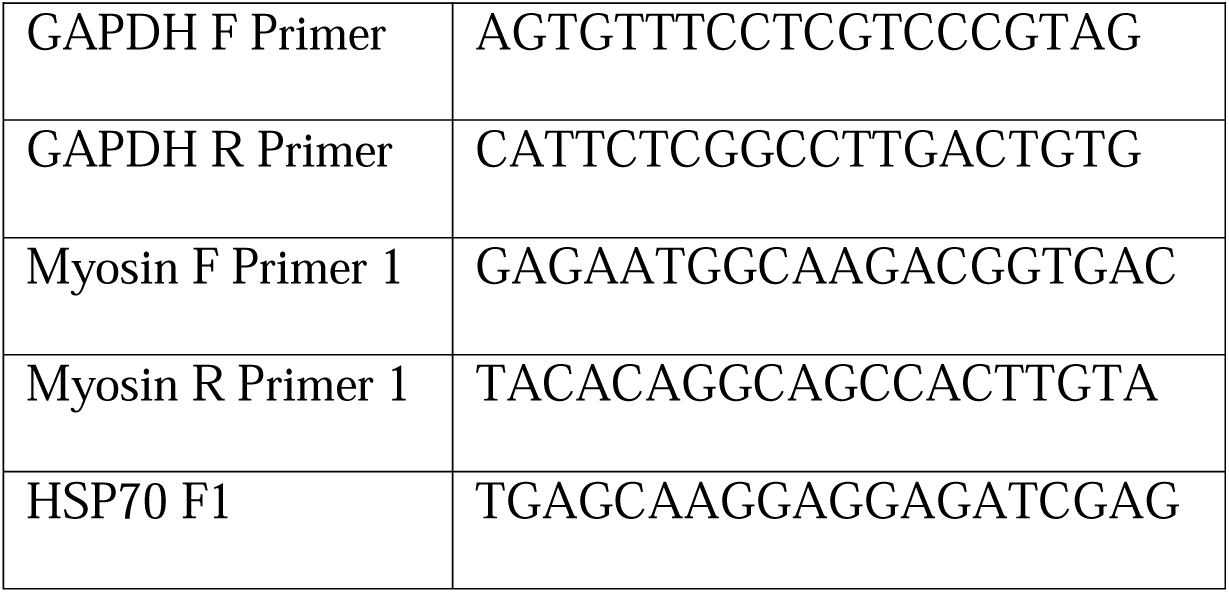
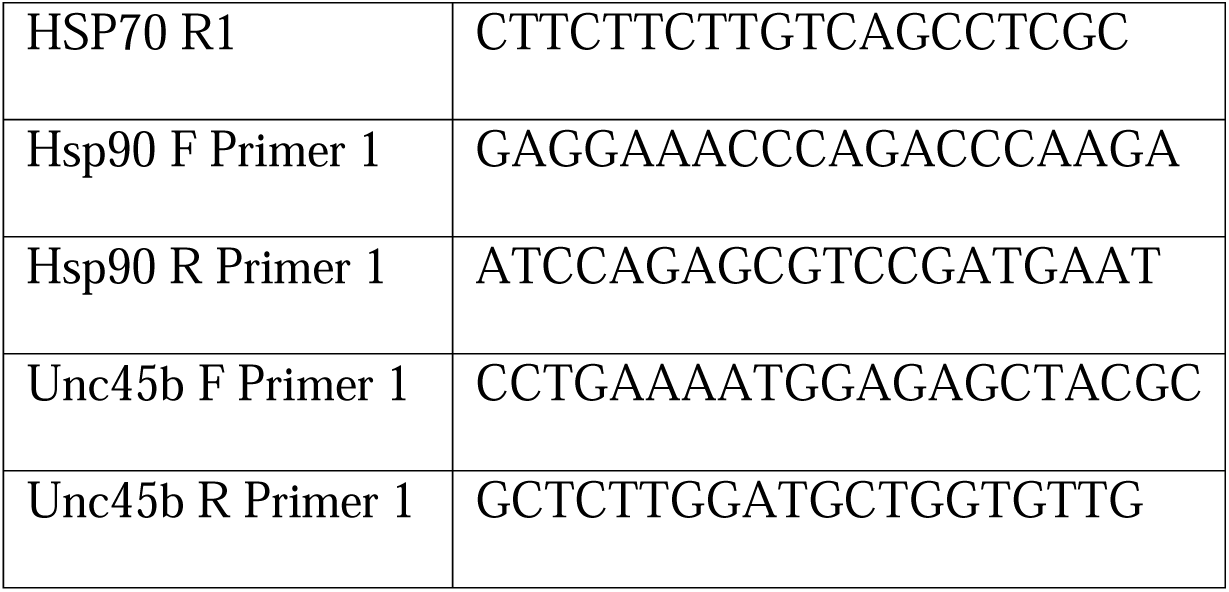

### 4.4. Immunofluorescence microscopy

Hearts were isolated, snap frozen in 2-methylbutane and embedded into cryogenic molds using OCT medium. 10 μm thick sections were obtained by cryosectioning using a cryostat, and the sections collected onto the surface of Colorfrost Plus (ThermoFisher) slides, dried, and stored at −80° until further use. We then used the acetone fixation and immunostaining protocol as described in Blondelle et al. (2019). Primary antibodies used: anti-α-actinin (Millipore A7811 mouse monoclonal at 1:800), anti-Hsp70 (Proteintech 10995-1-AP affinity purified rabbit polyclonal at 1:200), anti-Unc45b (Proteintech 21640-1-AP, affinitiy purified rabbit polyclonal at 1:200), and anti-N-cadherin (Santa Cruz sc-59987 mouse monoclonal at 1:100). The secondary antibody, used at 1:200 dilution, was anti-mouse Alexa 594, purchased from Invitrogen. Images for Figure 1B, C were captured with a Nikon A1R HD25 confocal system, and images for Figure 6 were captured with a Zeiss confocal system (LSM510) equipped with an Axiovert 100M microscope using Apochromatic 1.4 numerical aperture oil immersion objectives, x60 or x63, respectively. The color balances were adjusted using Adobe Photoshop (Adobe).

### 4.5. Statistics

Unpaired student t-test was used to determine statistical significance of differences. (https://www.graphpad.com/quickcalcs/ttest1/?format=sem).

## Abbreviations

Hsp: Heat shock protein
Unc: Uncoordinated

## ACKNOWLEDGEMENTS

PDB would like to thank Dr. Teresa Kee at the Case Western School of Medicine for providing some of the young and old mouse heart samples, and Drs. Pravin Dubey and Sarojini Singh in the Krishnamurthy lab for help and advice, Sarojini Singh for helping with mice dissection.

## Funding

This work was supported by the National Institutes of Health [grant numbers R01HL160693 and R01AG088215 to GMB], and Department of Defense [grant number HT9425-23-1-0278, PR220330 to PK]. PDB is supported by a Blazer Presidential Elite Scholarship, University of Alabama Birmingham. The funders had no role in the study design, data collection, and analysis, decision to publish, or manuscript preparation.

## CONFLICT OF INTEREST STATEMENT

None declared.

## AUTHOR CONTRIBUTIONS

Conceptualizations, P.D.B., P.K., H.Q., G.B.; methodology, P.D.B, A.N., G.B., L.S., J.K., formal analysis, P.D.B., A.N., G.B; investigation, P.D.B., A.N., L.S., G.B; data curation, P.D.B, A.N., L.S., G.B.; writing-original draft preparation, P.D.B., and G.B; writing-review and editing, P.D.B., G.B., P.K.; supervision, G.B., and P.K.; funding acquisition, G.B., and P.K. All authors have read and agreed to the published version of the manuscript.

## DATA AVAILABILITY STATEMENT

All data generated or analyzed during this study are included in this published article and its supplementary information files. Raw datasets and detailed protocols used during the current study are available from the corresponding author upon reasonable request.

## CONSENT TO PARTICIPATE

None applicable.

## CONSENT FOR PUBLICATION

All the authors have read and approved the final manuscript.

## Notes

### Competing Interest Statement

The authors have declared no competing interest.

### Summary of Updates

New data showing localization of HSP-70 and UNC-45b in cardiomyocytes. Removal of myosin staining. Corresponding changes to Results and Discussion.

## REFERENCES

Ao, W., and Pilgrim, D. (2000). Caenorhabditis elegans UNC-45 is a component of muscle thick filaments and colocalizes with myosin heavy chain B, but not myosin heavy chain A. J Cell Biol. 148, 375–384.

Banypersad SM, Moon JC, Whelan C, Hawkins PN, Wechalekar AD (2012-04-24). “Updates in Cardiac Amyloidosis: A Review“. Journal of the American Heart Association. 1 (2): e000364. doi:10.1161/JAHA.111.000364. ISSN 2047-9980. PMC 3487372. PMID 23130126.

Barral, J.M., Bauer, C.C., Ortiz, I., and Epstein, H.F. (1998). Unc-45 mutations in Caenorhabditis elegans implicate a CRO1/She4p-like domain in myosin assembly. J. Cell Biol. 143, 1215–1225.

Barral, J.M., Hutagalung, A.H., Brinker, A., Hartl, F.U., and Epstein, H.F. (2002). Role of the myosin assembly protein UNC-45 as a molecular chaperone for myosin. Science 295, 669–671.

Bhattacharya, K. Weidenauer, L., Luengo, T.M., Pieters, E.C., Echeverrria P.C., Bernasconi, L., Wider, D., Sadian, Y., Koopman, M.B., Villemin, M., Bauer, C., Rudiger, S.G.D., Qaudroni, M., and Picard, D. (2020). The Hsp70-Hsp90 co-chaperone Hop/Stip1 shits the proteostatic balance from folding towards degradation. Nature Comm. 11, 5975.

Blondelle, J., Marrocco, V., Clark, M., Desmond, P., Myers, S., Nguyen, J., Wright, M., Bremner, S., Pierantozzi, E., Ward, S., Esteve, E., Sorrentino, V., Ghassemian, M., and Lange, S. (2019). Murine obscurin and Obsl1 have functionally redundant roles in sarcolemmal integrity, sarcoplasmic reticulum organization and muscle metabolism. Commun. Biol. 2, 178.

Bujalowski, P.J., Nicholls, P., Garza, E., and Oberhauser, A.F. (2018). The central domain of UNC-45 chaperone inhibits the myosin power stroke. FEBS Open Bio. 8, 41–48.

Chen, D., Li, S., Singh, R., Soubettem S., Sedlmeier, R., and Epstein, H.F. (2012). Dual function of the UNC-45b chaperone with myosin and GATA4 in cardiac development. J. Cell Sci. 125, 3893–3903.

Dubey, P.K., Dubey, S., Singh, S., Bhat, P.D., Pogwizd, S., Krishnamurthy, P. (2024). Identification and development of Tetra-ARMS PCR-based screening test for a genetic variant of OLA1 (Tyr254Cys) in the human failing heart. PLoS ONE 19(6): e0293105. 10.1371/journal.pone.0293105

Dutta, S., Sengupta, P. (2016). Men and mice:relating their ages. Life Sci 152:244–248. https://doi.10.1016/j.lfs.2015.10.025

Epstein, H.F., and Thomson, J.N. (1974). Temperature-sensitive mutation affecting myofilament assembly in Caenorhabditis elegans. Nature 250, 579–580.

Etard, C., Roostalu, U., and Strahle, U. (2008). Shuttling of the chaperones Unc45b and Hsp90a between the A band and the Z line of the myofibril. J. Cell Biol. 180, 1163–1175.

Fikrle M, Paleček T, Kuchynka P, Němeček E, Bauerová L, Straub J, et al. (2013-02-01). “Cardiac amyloidosis: A comprehensive review“. Cor et Vasa. 55 (1): e60–e75. doi:10.1016/j.crvasa.2012.11.018. ISSN 0010-8650.

Gazda, L., Pokrzywa, W., Hellerschmied, D., Lowe, T., Forne, I., Mueller-Planitz, F., Hoppe, T., and Clausen, T. (2013). The myosin chaperone UNC-45 is organized in tandem modules to support myofilament formation in C. elegans. Cell 152, 183–195.

Gaziova, I., Moncrief, T., Christian, C.J., Villarreal, M., Powell, S., Lee, H., Qadota, H., White, M.A., Benian, G.M., and Oberhauser, A.F. (2020). Mutational analysis of the structure and function of the chaperoning domain of UNC-45B. Biophys. J. 119, 1–12.

Herndon, L.A., Schmeissner, P.J., Dudaronek, J.M., Brown, P.A., Listner, K.M., Sakano, Y., Paupard, M.C., Hall, D.H., and Driscoll, M. (2002). Stochastic and genetic factors influence tissue-specific decline in ageing C. elegans. Nature 419, 808–814.

Kachur, T.M., and Pilgrim, D.B. (2008). Myosin assembly, maintenance and degradation in muscle: Role of the chaperone UNC-45 in myosin thick filament dynamics. Int. J. Mol. Sci. 9, 1863–1875. 10.3390/ijms9091863.

Kaiser, C. M., Bujalowski, P. J., Ma, L., Anderson, J., Epstein, H.F., and Oberhauser, A.F. (2012). Tracking UNC-45 chaperone-myosin interaction with a titin mechanical reporter. Biophys. J. 102, 2212–2219.

Lee, C.F., Melkani, G.C., and Berstein, S.I. (2014). The UNC-45 mysoin chaperone: from worms to flies to vertebrates. Int. Rev. Cell Mol. Biol. 313, 103–144.

Matheny, C.J., Qadota, H., Bailey, A.O., Valdebenito-Silva, S., Oberhauser, A.F., and Benian, G.M. (2024). The myosin chaperone UNC-45 has an important role in maintaining the structure and function of muscle sarcomeres during adult aging. Mol. Biol. Cell 35:ar98, 1-18.

Melkani, G.C., Bodmer, R., Ocorr, K., and Bernstein, S.I. (2011). The UNC-45 chaperone is critical for establishing myosin-based myofibrillar organization and cardiac contractilitiy in the Drosophila heart model. PLoS ONE 6(7): e22579.

Morano, K. A. (2007). New tricks for an old dog: the evolving world of Hsp70. Annals of the New York Academy of Sciences. 1113 (1): 1–14. doi:10.1196/annals.1391.018. PMID 17513460. S2CID 20917046.

Price, M.G., Landsverk, M.L., Barral, J.M., Epstein, H.F. (2002). Two mammalian UNC-45 isoforms are related to distinct cytoskeletal and muscle-specific functions. J. Cell Sci. 115, 4013–4023.

Sheydina, A., Riordon, D.R., Boheler, K.R. (2011). Molecular mechanisms of cardiomyocyte aging. Clin. Sci. 121, 315–329.

Solomon, V., and Goldberg, A.L. (1996). Importance of the ATP-ubiquitin-proteasome pathway in the degradation of soluble and myofibrillar proteins in rabbit muscle extracts. J. Biol. Chem. 271, 26690–26697. 10.1074/jbc.271.43.26690.

Tavaria, M., Gabriele, T., Kola, I., Anderson, R. L. (1996). A hitchhiker’s guide to the human Hsp70 family. Cell Stress & Chaperones. 1 (1): 23–28. doi:10.1379/1466-1268(1996)001<0023:ahsgtt>2.3.co;2. PMC 313013. PMID 9222585

Venolia, L., Ao, W., Kim, S., Kim, C., and Pilgrim, D. (1999). unc-45 gene of C. elegans encodes a muscle-specific tetratricopeptide repeat-containing protein. Cell Motil. Cytoskeleton. 42, 163–177.

Vijayakumar, A. Wang, M., and Kailasam, S. (2024). The senescent heart—“age doth wither its infinite variety”. Int. J. Mol. Sci. 25, 3581.

